# Globular domain histone H3R131C mutation remodels chromatin accessibility to promote oncogenic transcriptional programs

**DOI:** 10.64898/2026.05.27.728333

**Authors:** SV Lohano, K Sad, AJ Kyle, E Hill, SA Sloan, AH Corbett, JM Spangle

**Author notes:** Authors contributed equally (SV Lohano, K Sad).

## Abstract

Histone mutations, characterized as oncohistones, have emerged as important oncogenic driver events by altering chromatin structure and/or chromatin modifications, thereby dysregulating gene expression. While H3 tail domain oncohistone mutations such as H3K27M and H3K36M are well characterized, it is currently unknown whether mutations within the H3 globular domain represent oncogenic driver events. Using publicly available cancer patient tumor data, here we identify H3R131C as a recurrent histone H3 globular domain mutation. H3R131C mutation is present in diverse human tumors including breast and bladder cancers. Phenotypic assays demonstrate that H3R131C expression does not augment cellular proliferation but enhances cellular migration and invasion. H3R131C frequently co-occurs with mutations in oncogenes including PIK3CA and ESR1, and tumor suppressors TP53 and CDKN2A with an allele frequency consistent with H3R131C representing a subclonal event that enhances tumor fitness rather than initiating transformation. Mechanistically, ATAC-seq reveals that H3R131C expression increases chromatin accessibility, with enrichment of AP-1/bZIP transcription factor motifs at gained accessible regions. Integration of chromatin accessibility and transcriptomic profiling identifies concordant upregulation of pro-oncogenic target genes including PGF, SOX5, and CCDC88C, supporting a model in which H3R131C destabilizes the nucleosome to remodel chromatin and activate transcriptional programs associated with cellular migration, angiogenesis, and epithelial plasticity. Collectively, these findings identify H3R131C as a functionally active globular domain oncohistone that reshapes the epigenome to promote oncogenic gene expression.

## Introduction

Cancer is a complex disease that arises from genomic alterations that change cellular processes, including uncontrolled cell proliferation, DNA damage response, and the dysregulation of gene expression (1-3). DNA in eukaryotic cells is packaged into chromatin, which is organized into repeating nucleosome subunits. Each nucleosome is comprised of 146-147bp of DNA, which wraps around a hetero-octamer of the histone proteins H2A, H2B, H3, and H4 (4, 5)..Histones are the structural proteins that package DNA into chromatin, with each histone core protein containing a flexible N-terminal tail that extends beyond nucleosome core, consisting of histone globular domains and DNA. Beyond their structural role, histones are critical regulators of gene expression and genome stability, influencing processes such as transcription, replication, and DNA damage repair through dynamic chromatin remodeling and post-translational modifications (PTMs) (1, 6). These histone PTMs include methylation, acetylation, SUMOylation, phosphorylation, ubiquitination, acylation, lactylation, and many others (7-9). Moreover, the protein machinery that reads, writes, and erases histone PTMs are altered in cancer and can serve as oncogenic driver event that and promote tumorigenesis and influence patient prognosis and therapeutic response (10, 11).

Recurrent mutations in histone genes have been identified across a variety of human cancers. The identification of recurrent histone mutations that drive oncogenic activity in *in vitro* and *in vivo* tumor models has led to the term, “oncohistone” (12). While growing in number, characterized histone H3 oncohistone mutations including H3K27M, H3K36M, and H3G34R/V/W/L occur in a variety of human cancers including pediatric diffuse intrinsic pontine glioma (DIPG) (H3H27M), chondroblastoma and squamous cell cancer of the head and neck (SCCHN; H3K36M) (13, 14), pediatric glioblastomas (H3G34V/R), and giant cell tumors of the bone (H3.3G34W/L) (15). These genomic alterations lead to the production of histone H3 proteins with a mutation in the histone H3 N-terminal tail domain and alters the ability of the histone protein to support PTMs (16, 17). Because oncohistone-driven cancers are associated with poor patient survival, it is important to study how and why histone mutations support tumorigenesis. Collectively, published studies highlight the oncogenic potential of a small subset of identified genomic alterations in histone genes. To date, genomic alterations that lead to changes in more than 20 H3 amino acids have been defined in patient tumors, the majority of which remain uncharacterized (18).

While the mechanisms by which oncohistone proteins drive cancer varies, prior studies suggest that these mutations alter nucleosome integrity (19), chromatin accessibility (20), and/or interfere with histone PTMs (16) (21). The H3K27M oncohistone binds EZH2, the catalytic subunit of Polycomb Repressive Complex 2 (PRC2), thereby impairing EZH2 methyltransferase activity on histone substrates, which results in the non-uniform reduction of repressive H3K27 trimethylation (H3K27me3) and drives enhanced gene expression (22). A structural study reveals that the inhibitory effect of H3K27M on PRC2 activity is comparable to small-molecule EZH2 inhibitors, indicating that the mutation likely acts to block EZH2 activity by disrupting substrate binding to the catalytic SET domain of EZH2, through PRC2-independent effects (23). The H3K36M oncohistone also binds the H3K36 methyltransferases NSD1/2 and SETD2, decreasing H3K36me2/me3. A loss of H3K36 methylation is accompanied by the redistribution of PRC2-catalyzed H3K27me3 and thus reprogramming of the epigenetic landscape (24). In contrast to N-terminal tail alterations, mutations in histone globular domains can promote cancer via distinct mechanisms that are at least partially independent of changes to the histone PTM landscape. The H2B oncohistone H2BE76K changes the charge in the H2B globular domain, creating electrostatic repulsion with coordinating amino acids such as H4R92, which leads to a conformational change in the α3-helix of H4. The change in conformation disrupts the salt bridge between H2BE76-H4R92 and destabilizes the nucleosome(19, 25, 26).

While the mechanisms driving oncogenic activity vary amongst histone driver mutations, in all cases gene expression is re-wired, which may present opportunities for therapeutic targeting. In the case of H3K27M-mutant DIPGs, transcription of the dopamine receptors DRD2/3 are upregulated. To date, the DRD2/3 small molecule antagonist ONC201 is the first and only FDA approved clinical therapy for the treatment of H3K27M-mutant diffuse midline glioma. While H3K27M-mutant diffuse midline glioma patients treated with ONC201 exhibit therapeutic benefit with a median overall survival benefit of 22 months in ONC201 treated patients compared to 12 months for standard of care, additional therapeutic approaches for the treatment of oncohistone-driven cancers are in preclinical and clinical investigation and include immunotherapeutic approaches (27, 28) (29, 30). Preclinical and clinical studies focused on known oncohistones underscore the importance of histone mutation in biological, functional, and clinical contexts; however, these studies also illustrate critical gaps in our knowledge of how histone mutations shape functional biology outcomes to contribute to cancer initiation, progression, and therapeutic response.

Here we provide the initial characterization of the genomic alterations in histone H3 genes that result in the expression of the globular domain H3R131C mutant protein. While these genomic alterations have been defined in multiple pan-cancer studies and histone mutation datasets as a recurrent alteration (12, 19, 31), the functional consequences of H3R131C expression are currently unknown. Here we leverage publicly available patient tumor data (COSMIC and cBioPortal) to identify H3R131C as a recurrent globular domain mutation found in diverse solid tumors and hematological malignancies including breast cancer. We engineer models of breast specific H3R131C expression and find that while stable expression of H3R131C increases cell migration and invasion, H3R131C does not increase cell proliferation and clonogenic growth. Mechanistic studies suggest that H3R131C expression increases chromatin accessibility and gene expression, leading to a more permissive chromatin state that is characterized by a gain in differentially accessible regions and associated with upregulation of key transcriptional programs associated with migration and epithelial plasticity. Collectively, these studies support H3R131C alteration as a probable oncogenic event and provides mechanistic insight into how chromatin state may influence gene expression to alter transcriptional output to drive oncogenic phenotypes in models that express H3R131C.

## Results

### Genomic alterations that change histone H3 R131 recur in human cancers

While H3 tail domain mutations such as H3K27M and H3K36M/R are well-established oncogenic drivers that perturb histone PTMs in cis and/or in trans to dysregulate transcription, whether mutations within the histone H3 globular domain similarly drive cancer remains poorly understood. To determine whether genomic alterations that change the histone H3 globular domain amino acids are associated with human cancers, we surveyed the COSMIC (32) and cBioPortal (33) publicly available adult human cancer databases and identified recurrent mutations that alter H3R131. Among the H3R131 alterations, mutations that result in the change of arginine 131 to that of a cystine (H3R131C), are the most common, found in 27 patient tumors (**Figure 1A, Figure S1A**). While multiple histone H3 genes harbor genomic alterations at R131, mutations are the most common in genes encoding H3.1 (**Figure 1B**). While H3R131 alterations occur across cancer types such as breast, bladder, lung, and skin, which is similar to the distribution described for the known oncohistone H2BE76K (19, 25, 26) (**Figure S1B**), H3R131C occurs most frequently in bladder and breast cancers, as well as melanoma (**Figure 1C**). Localized to the globular domain of histone H3 (**Figure 1D**), H3R131 is not surface accessible within the nucleosome. Structural modeling reveals that H3R131 occupies the central core of the H3 globular domain (**Figure 1E**). Because alteration of basic and polar arginine to uncharged cysteine may disrupt local interactions within the histone and/or nucleosome, these data suggest that H3R131C mutation could impact nucleosome stability via histone-histone interactions.

**Figure 1:**
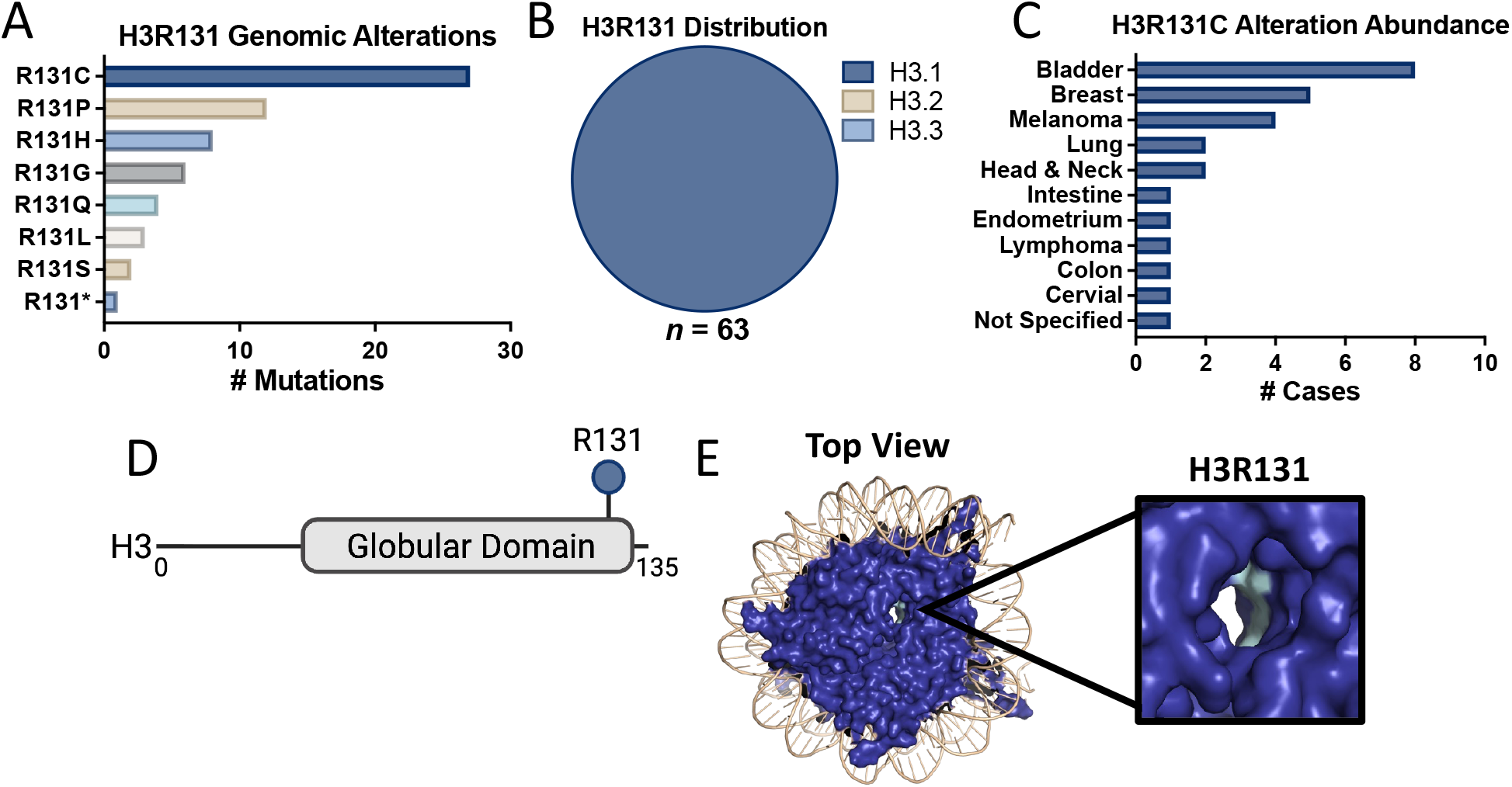
Histone H3R131 alterations recur in human cancers. Data acquired from cBioPortal and COSMIC databases. A) Number of genomic alterations defined that alter histone H3R131 in adult tumors. B) Distribution of histone H3 genes that harbor genomic alterations that change H3R131. C) Distribution of identified cancer diagnoses that harbor changes in H3R131C. D) Schematic diagram of histone H3 proteins with R131 annotated. E) *In silico* modeling representing top view (left) and magnified view of the nucleosome core. DNA = tan; H3R131 = light blue; remainder of nucleosome proteins H2A, H2B, H3 and H4 = blue. PDB ID 5X7X.

To begin to address the possible cancer driver nature of the H3R131 alteration, we utilized a linear regression model to delineate the probability that H3R131 genomic alteration is associated with a cancer diagnosis. Compared to the bona-fide H3 oncohistones H3K27M and H3K36M, H3R131 alteration is associated with a lower probability of cancer diagnosis (H3R131: 0.677, 95% CI [0.576, 0.764]; H3K27M: 0.976, 95% CI [0.963, 0.984]; H3K36M: 0.855, 95% CI [0.792, 0.902, respectively), while predominantly associating with a cancer diagnosis. In contrast, alteration of a random H3 residue, H3R63, is not associated with a cancer diagnosis (**Figure 2A**). Genomic datasets from a diverse and non-diseased population, GnomAD (34), highlight that similar to H3K27 and H3K36 alterations, H3R131 is uncommon in the random non-diseased population (**Figure 2B**), supporting the conclusion that H3R131 alterations are unlikely to be general SNPs in the human population. We also find that H3R131 allele frequency captured in patient tumors is 0.236, compared to H3K27 (0.368) and H3K36 (0.284) (**Figure 2C**). While the diagnosed cancer allele frequency for H3R131 is not significantly different than that of H3K27 or H3K36, these data support the conclusion that H3R131 genomic alteration is likely a subclonal event. In support of a subclonal event, aneuploidy scores in patient tumor samples harboring the H3R131C alteration is 7.28 compared to those harboring H3K27 (11.25) or H3K36 (6.85) alterations (**Figure 2D)**, which is consistent with H3R131C arising in the context of broader chromosomal instability. Further, analysis of patient genomic data demonstrates that H3R131C alterations co-occur with oncogenic driver genomic alterations including *PIK3CA, ESR1*, and *FGFR3*, and/or alterations in tumor suppressor genes, including *TP53, CDKN2A*, and *NF1* (**Figure 2E**).

**Figure 2:**
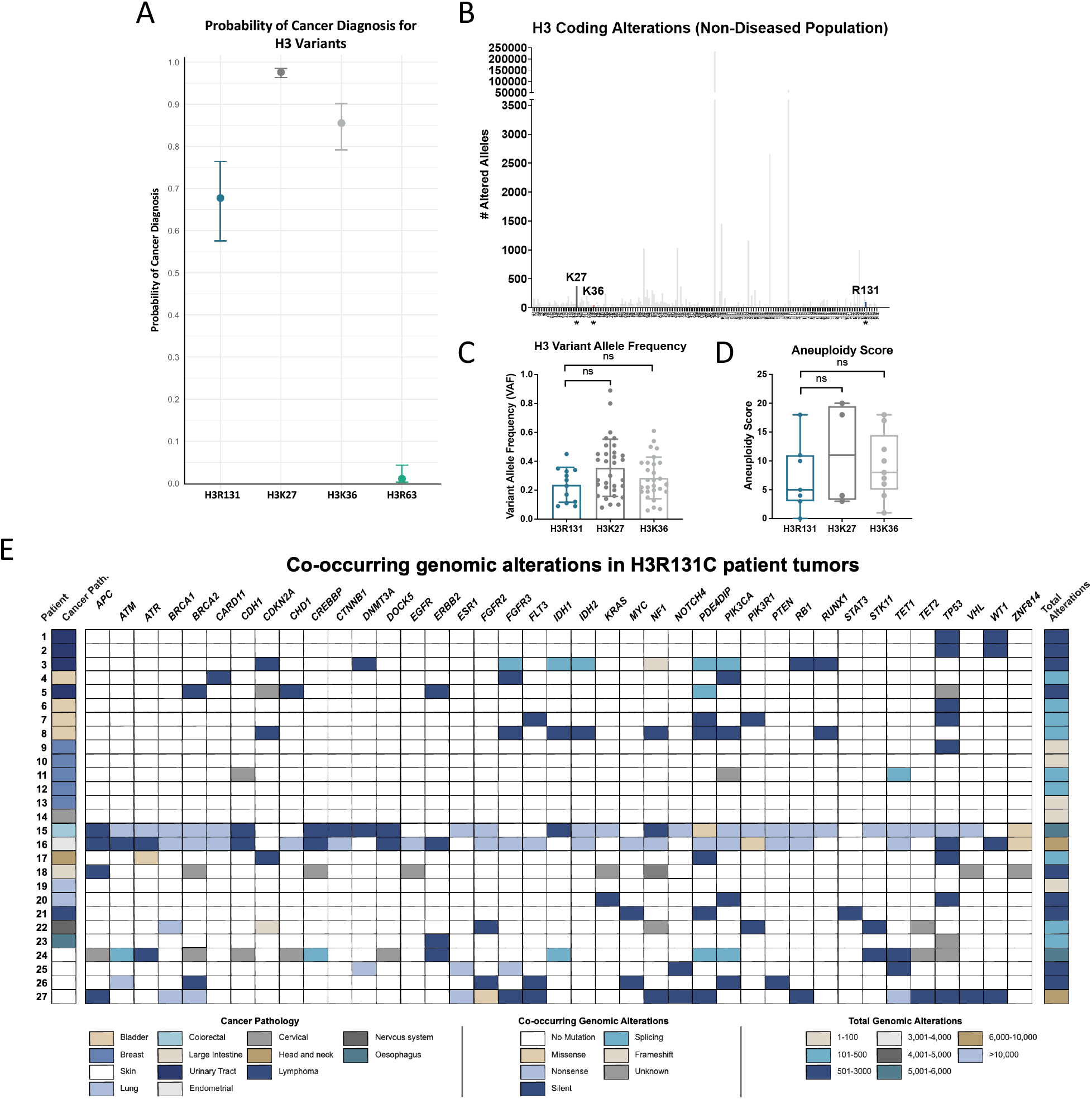
Alteration of H3R131 is associated with human cancer and co-occurs with common oncogenic driver events. A) Logistic regression model demonstrating the probability of cancer diagnosis associated with H3 mutations, where 0 = no diagnosis and 1 = cancer diagnosis. B) Frequency of genomic alterations that encode H3R131 changes defined in the large, non-diseased GnomAD human genomic database. C) Variant alle frequency for H3R131, compared to the known oncohistones H3K27, and H3K36, as reported in COSMIC and cBioPortal. One way ANOVA was performed for statistical significance. D) Boxplots showing aneuploidy score for patient samples reported in cBioPortal. One way ANOVA was performed for statistical significance. E) Overview of cancer pathology, ploidy score, co-occurring genomic alterations in common oncogenes and tumor suppressors, and total number of genomic alterations in patient tumors characterized by H3R131C mutant expression.

### H3R131C does not augment cellular proliferation

In order to examine the cellular impact of H3R131C expression, we engineered an *in vitro* model to study the role of H3R131C in breast transformation and oncogenic activity, because H3R131C alterations recur in breast cancer (**Figure 1C**). We engineered hTERT immortalized but untransformed human mammary epithelial cells (HMECs) that express a dominant negative p53 mutant protein (DD) to stably overexpress wildtype H3 (H3) or H3R131C with a C-terminal TY1 epitope tag. HMECDD cells were selected because the expression of dominant negative p53 in this context requires an additional oncogenic driver event (e.g., *KRAS*^*G12D*^, *PIK3CA*^*E542K/E545K/H1047R*^, or others) to produce a transformed phenotype (35-38). The HMECDD cell line was transduced with retroviral vectors expressing a C-terminal TY1 epitope-tagged H3 or H3R131C. While the cells overexpress ectopic H3 (**Figure 3A, Figure S2A**), quantitation demonstrates that the overexpression of H3.3 represents a small minority of total H3 expressed in the cells; approximately 7.95% of total H3 (**Figure 3B**) and 0.93% of total H3.3 in HMECDD cells (**Figures 3B and 3C**). These results are consistent with the ectopic overexpression used in other oncohistone studies. (19, 26, 35, 39, 40).

**Figure 3:**
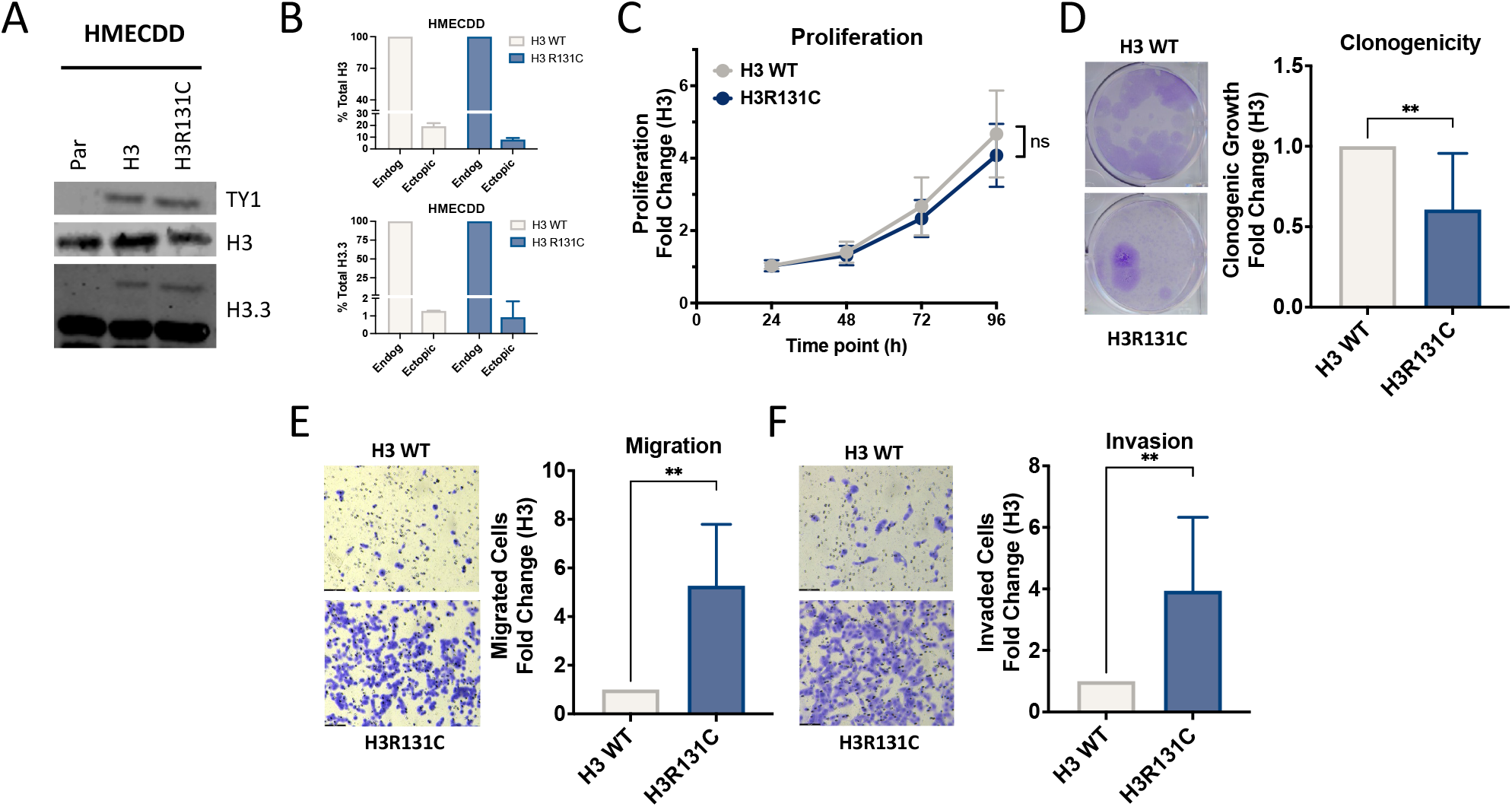
H3R131C expression supports cellular phenotypes associated with transformation. A) HMECDD cells stably transduced with pBabePuro H3.3-TY1 or H3.3R131C and lysates acid extracted. Lysates were immunoblotted with the indicated antibodies. Representative images shown, *n= 3*. B) Quantification of (A). Percent of total H3 (top) and total H3.3 (bottom) expression that represents ectopic total H3 and total H3.3 for HMECDD cells. C) Stable HMECDD cells expressing the indicated H3 mutant proteins were seeded and cell proliferation measured over the indicated time course, *n = 3*. D) HMECDD cells stably transduced with the indicated H3 plasmids were seeded and cell clonogenicity measured after 21 d. Representative images shown on the left, and quantification on the right, *n = 3*. E) HMECDD cells stably transduced with the indicated H3 plasmids were seeded and cell migration through an 8.0 μm filter was measured after 8 hr, representative images shown on left and quantification on right, *n = 3*. F) HMECDD cells stably transduced with the indicated H3 plasmids were seeded and cell invasion through an 8.0 μm filter coated with 1 mg/ml matrigel was measured after 48 hr. Representative images shown on the left and quantification on the right, *n = 3*.

We then tested whether expression of H3R131C augments cellular proliferation. Expression of H3R131C does not change the cellular proliferation rate compared to H3 overexpressing HMECDD cells in proliferation assays (**Figure 3D**). To more quantitatively examine cellular proliferation, we utilized the Incucyte platform to measure population doublings and found that H3R131C does not increase cellular proliferation compared to H3 (**Figure S2B**). We then examined whether H3R131C expression could support robust growth when cultured in limited dilution. Clonogenic assays also suggest that H3R131C expression does not alter the ability of HMECDD cells to form colonies when grown in limited dilution, when compared to H3 (**Figure 3E**). Taken together, these *in vitro* results suggest that H3R131C may not be involved in cellular processes associated with cancer initiation, as H3R131C expression does not provide cells with a proliferative advantage. These data may reflect the H3R131 allele frequency in **Figure 2**, highlighting the possibility that H3R131 alteration is clonal and is unlikely the initial cancer driver in this tumor context, but rather represents a genomic alteration that enhances the tumor fitness (41).

### H3R131C enhances cellular phenotypes associated with advanced stage disease

To examine whether H3R131C expression may augment cellular properties associated with migration and invasion, which function as surrogate *in vitro* markers of metastasis, we examined cellular migration and invasion via boyden chamber assay. Expression of HMECDD H3R131C significantly increases Boyden chamber-associated migration compared to HMECDD cells that express wildtype H3 (**Figure 3F**). Moreover, expression of HMECDD H3R131C enhances invasion through matrigel-coated Boyden chambers in comparison to HMECDD H3 cells (**Figure 3G**). Collectively, these data suggest that H3R131C may play a role in cellular functions that are associated with advanced stage disease, such as metastasis.

### H3R131C perturbs the chromatin landscape

To understand the mechanistic basis behind H3R131C-associated oncogenic phenotypes, we considered the structural and biochemical properties of H3R131 compared to H3C131, and how this biochemical change may alter the function of the nucleosome. *In silico* modeling suggests that H3R131 maintains a stable nucleosome core by mediating interactions with surrounding H3 residues H3D106 and H3E133. Change to H3C131 alters the amino acid charge, creating a repulsive force between H3C131 and both H3D106 and H3E133 (**Figure 4A**). Collectively, these data support the hypothesis that H3R131C expression may lead to a reduction in nucleosome stability and/or an increase in chromatin accessibility.

**Figure 4:**
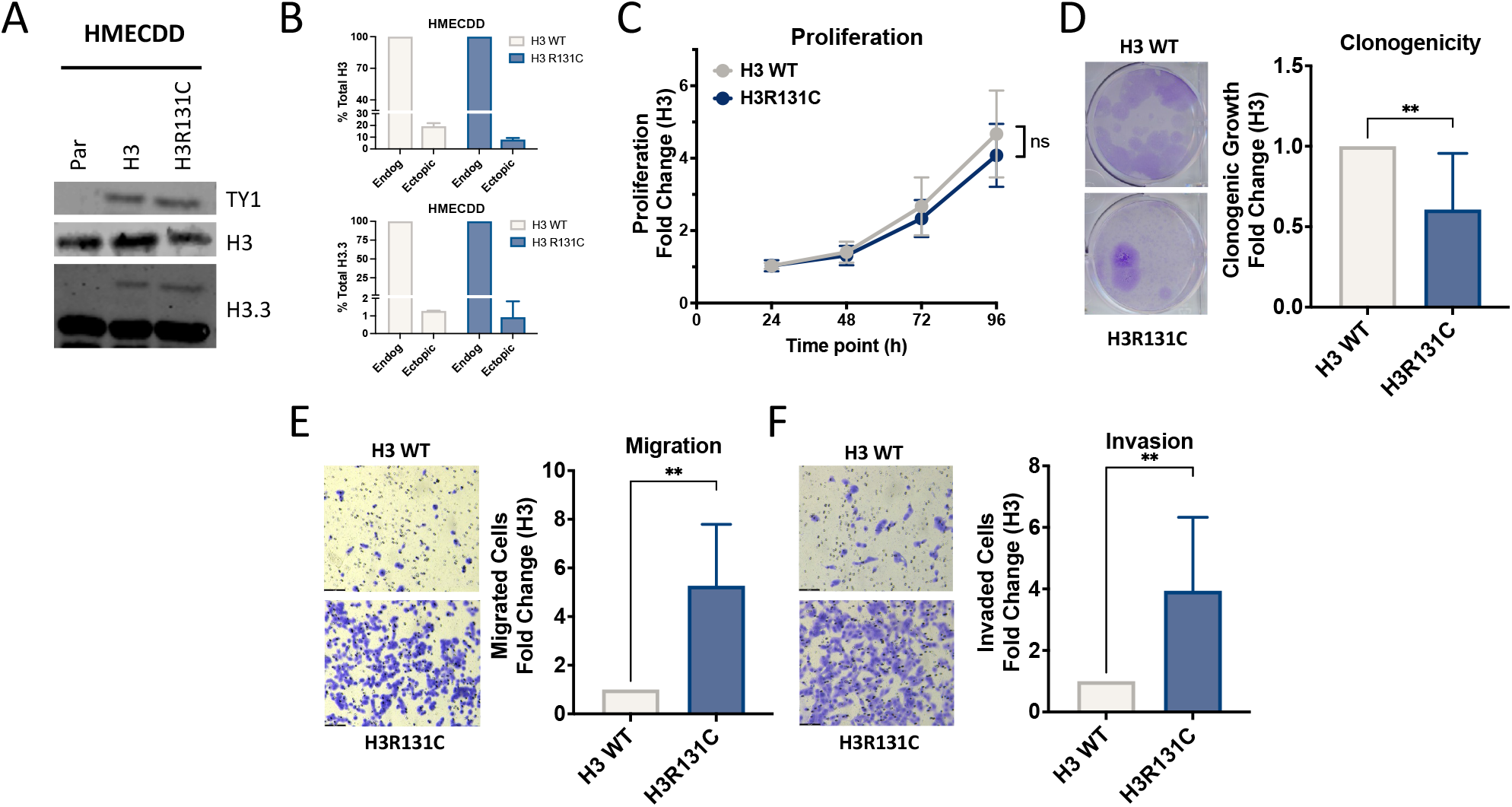
H3R131 supports an open chromatin state that drives changes in cellular function. A) *In silico* modeling of the intramolecular interactions in the wildtype nucleosome or the nucleosome containing H3R131C, using PDB 5X7X. B) Volcano plots depicting genomic loci with increased and decreased accessibility in HMECDD cells stably expressing H3R131C compared to H3 (total n = 1652 loci; dashed lines indicate log2 fold change cutoff of +/-1.5) C) Tornado plots assessing chromatin accessibility in HMECDD cells stably transduced with H3 or H3R131C. Representative plot showing the top 15,000 peaks in each cell line from a single biological replicate, *n = 2*. D) Genomic distribution of the differentially accessible regions between HMECDD cells expressing H3R131C or H3. E) Tornado plot representing loci that exhibit gained accessibility (top) or lost accessibility (bottom) in H3R131C expressing HMECDD cells compared to H3. F) HOMER motif enrichment analysis from ATAC-seq datasets. G) GO depicting upregulated biological pathways from HMECDD H3R131C-TY1 cells compared to HMECDD H3-TY1 cells.

To determine whether changes to chromatin accessibility may contribute to the observed oncogenic phenotypes (**Figure 3**), we performed Assay for Transposase-Accessible Chromatin using sequencing (ATAC-seq) (**Figure 4B, Figure S3A**). While H3R131C expression results in genomic regions characterized by both reduced (total of 92 decreased accessible loci) and increased accessibility (total of 437 loci with increased accessibility) compared to H3, H3R131C expression results in a more open chromatin conformation (**Figure 4C, Figure S3B**). To characterize the genomic distribution of these differentially accessible regions (DARs), we assessed their annotation relative to known genomic features. Regions of increased accessibility in H3R131C expressing cells are enriched at promoters and intronic regions, while regions of decreased accessibility are more broadly distributed across intronic and intergenic loci. (**Figure 4D**). Tornado plots stratified by gained and lost accessibility further confirm that H3R131C-associated chromatin remodeling is characterized predominantly by a gain of accessibility at a greater number of loci relative to those exhibiting reduced accessibility, suggesting a shift towards a more open chromatin state at specific genomic loci (**Figure 4E**). To identify transcription factors (TF) whose binding may be altered by H3R131C-driven chromatin remodeling, we performed TF motif enrichment analysis on the DARs using HOMER(42). H3R131C expression increases the accessibility of AP-1/bZIP transcription factor family motifs, as well as Fos, JunB, Fra1, AP-1, ATF3 TF motifs. These results suggest a role of H3R131C mutation in promoting oncogenic transcriptional programs driven by the AP-1 family of TFs (**Figure 4F**). We next mapped the DARs to the nearest gene using Cluster profiler and performed gene ontology enrichment analysis (GO). H3R131C increased accessible loci corresponds to signatures of biological processes that are involved in cell size, actin-mediated polymerization and projection, as well as insulin signaling (**Figure 4G**). Increased chromatin accessibility at loci associated with actin cytoskeletal organization has been linked to transcriptional reprogramming that promotes mesenchymal gene expression and enhanced migratory and invasive capacity in cancer cells (10), which is consistent with an observed H3R131C-driven increase in cellular migration and invasion (**Figure 2**). In contrast, GO enrichment does not identify any dysregulated biological processes from the loci characterized with decreased accessibility in H3R131C cells when compared to wildtype H3. Examination of GO molecular function and GO cellular component identifies increased chromatin accessibility signatures linked to other cellular functions including GTPase activity, cell leading edge, and cell ruffling, which also have known connections to cellular migration (43, 44)(**Figures S3C, S3D**)

### Chromatin conformation underlies H3R131C-associated transcriptional changes

Because changes in chromatin conformation can be associated with transcriptional state, we performed RNA sequencing in HMECDD cells that stably express H3R131C or H3. RNA-seq identified 94 significantly upregulated and 45 significantly downregulated genes in HMEC DD cells expressing H3R131C when compared to H3 (**Figure 5A, Figure S4A**). Fast gene set enrichment analysis (FGSEA) identifies concerted regulation of cancer-associated pathways by H3R131C expression: H3R131C expression upregulates gene signatures associated with hypoxia and MYC targets, while downregulating pathways associated with TGFβ and TNFα signaling (**Figure 5B**). Integration of the ATAC-seq and RNA-seq datasets suggests that stable H3R131C expression drives concordant regulation of a subset of genomic loci accessibility and transcript abundance (**Figure 5C**). An increase in genomic loci accessibility when mapped to the nearest gene, and concurrent increase in transcript abundance support the hypothesis that H3R131C-associated changes to chromatin accessibility drive concordant changes to transcription abundance, which may influence the biological function of cells expressing H3R131C.

**Figure 5:**
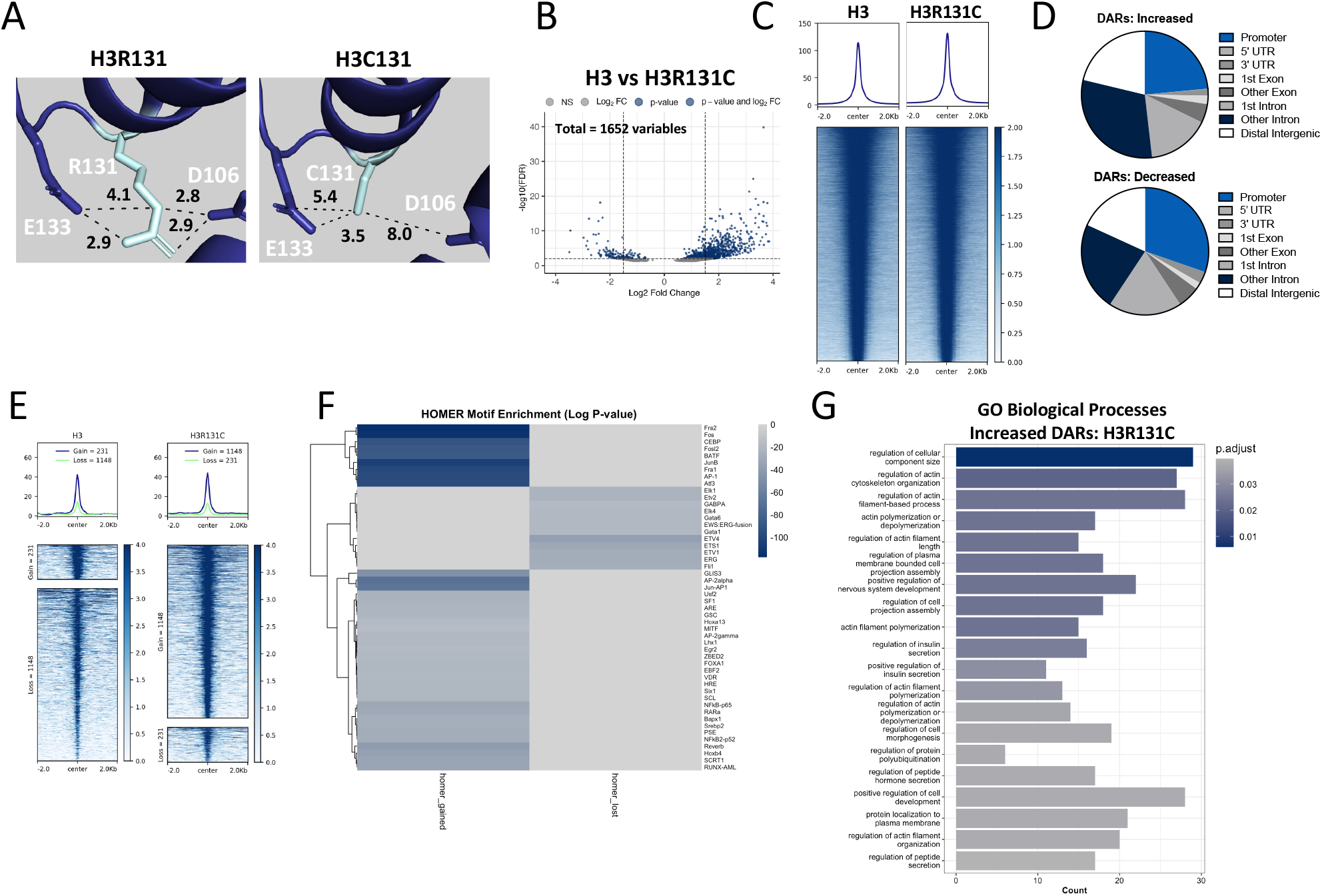
Chromatin accessibility and transcriptional changes underlie H3R131C oncogenic phenotypes. A) Volcano plot depicting increased and decreased transcript abundance in HMECDD cells stably expressing H3R131C compared to H3 where *P* ≤ .05 (blue) and log_2_fold change ≥ 1.5 (red).B) FGSEA depicting upregulated or downregulated hallmark pathways from HMECDD H3R131C-TY1 cells compared to HMECDD H3-TY1 cells (blue padj < 0.05, red padj > 0.05). C) Concordant analyses of differentially accessible loci and differentially expressed transcripts with fold change ≥ 1. Scatter plots display the relationship between ATAC-seq log2 fold change and RNA-seq log2 fold change. ™log10(ATAC adjusted p-value × RNA adjusted p-value) was used for statistical significance. D) Euler plots highlighting concordant data derived from the ATAC-seq and RNA-seq datasets. Left= shared increased accessible loci mapped to the nearest gene and increased transcript abundance. Right = shared decreased loci accessibility mapped to the nearest gene and decreased transcript abundance. E) Normalized read counts of indicated genes that are differentially expressed in HMECDD H3R131C-TY1 compared to H3-TY1 cells. F) IGV tracks from ATAC-seq and RNA-seq of selected genes PGF, SOX5, and CCDC88C.

### H3R131C remodels the genome to support expression of key oncogenic regulators

Examining the concordant regulation of chromatin accessibility and transcript abundance identifies a total of eight genes that are characterized by both increased accessibility and increased transcript abundance (CCDC88C, PGF, SOX5, SYNE2, ABCC2, SGPP2, RCN3, and GJB2) and a total of two genes characterized by decreased accessibility and decreased transcript abundance (GALNT5 and DRC11) (**Figure 5D**). Normalized read counts derived from our RNA-seq dataset highlights the increased transcript abundance of PGF, SYNE2, GJB2, and CCDC88C, and the decreased transcript abundance of GALNT5 and DRC11 in H3R131C expressing cells as compared to H3 (**Figure 5E**). Examination of the genomic ATAC- and RNA-sequencing tracks of the PGF, a secreted angiogenic factor and member of the vascular endothelial growth factor (VEGF) family, and SOX5, a transcription factor implicated in the maintenance of stem-like and mesenchymal cell states demonstrates an increase in chromatin accessibility, and an increase in RNA transcript abundance in HMECDD cells that stably express H3R131C in comparison to H3 (**Figure 5F, Figure S5**). Taken together, concordant accessible genomic loci/increased transcripts are consistent with the GO enrichment signatures (**Figure 4G, Figures S3C, S3D**) and support the hypothesis that H3R131C promotes an oncogenic gene expression program characterized by increasing transcripts associated with migration, angiogenesis, and cell plasticity. Collectively, these findings establish H3R131C as a potential oncohistone which can reshape the chromatin to regulate gene expression to support oncogenic transformation.

## Discussion

Here we identify genomic alterations that lead to the expression of histone H3R131C as recurrent in human cancers including breast. Our results suggest that H3R131C drives increased cellular migration and invasion, which are molecular properties typically associated with advanced stage cancer. Unlike the H3K27M and H3K36M oncohistones that occur within the N-terminal tail and perturb native histone posttranslational modifications (17) (16), H3R131C is embedded in the H3 globular domain where it stabilizes the nucleosome core structure through electrostatic interactions with neighboring residues H3D106 and H3E131. H3R131C expression is associated with increase in chromatin accessibility and enrichment of TF motifs associated with the AP-1 family of TFs. Transcriptomics suggest activation of gene signatures associated with hypoxia and MYC targets. Integration of our chromatin accessibility and transcriptomic data identifies concordant increased assessibility and increased transcript abundance of PGF, which is a pro-angiogenic secreted member of the VEGF family that promotes tumor cell motility and invasion (45) and the TF Sox5 (46), which together support oncogenic signatures associated with EMT and other functions associated with cancer. Taken together, our data support a model in which H3R131C destabilizes local nucleosome architecture, increases chromatin accessibility at discrete genomic loci such as PGF and SOX5, and drives transcriptional changes that support oncogenic progression particularly phenotypes associated with cellular migration, invasion, and metastasis.

A notable feature of H3R131C is that its expression in untransformed breast cells does not confer increased proliferation *in vitro*, which prompted us to consider how H3R131C may contribute to oncogenesis in the absence of a proliferative advantage. While H3K36M expression in mesenchymal progenitor cells is sufficient to induce sarcoma formation *in vivo* (17), our recent studies characterizing the globular domain oncohistone H3E50K suggest that H3E50K expression modestly increases cell proliferation and clonogenic growth, while considerably enhancing migratory and invasive potential in both immortalized but untransformed models and in the context of co-occurring *BRAF* mutations. These data suggest that globular domain oncohistones may be predisposed to drive phenotypes associated with cancer progression rather than initiation (35). This observation is consistent with the subclonal allele frequency of H3R131C (0.22) observed in patient tumors, and its co-occurrence with bona-fide oncogenic drivers including PIK3CA^E542K/E545K/H1047R^, as well as loss-of-function alterations in tumor suppressors such as TP53. Collectively, these data support a model in which H3R131C is not likely driving cancer initiation, but rather collaborates with known oncogenic drivers to support or accelerate biological processes associated with advanced stage disease. While more experiments are required to test this hypothesis, a possible explanation for these results may be the differences in mechanisms by which N-terminal tail H3 oncohistones function to drive oncogenic activity in comparison to H3 globular domain mutations: established N-terminal H3 oncohistones alter histone PTM patterns primarily in a dominant nature, and our studies suggest that H3 globular domain mutations alter chromatin accessibility and/or stability.

Histone H3R131 forms stabilizing electrostatic contacts with H3D106 and H3E133 within the hydrophobic core of the H3 globular domain; substitution of the positively charged arginine with uncharged cysteine is predicted to introduce repulsive forces that locally destabilize the nucleosome. Our ATAC-seq data (**Figure 4**) supports this model, demonstrating that H3R131C expression results in a net increase in chromatin accessibility, with 437 loci gaining and 92 loci losing accessibility relative to wildtype H3. This approximately 4.7-fold gain in accessible regions may suggest H3R131C may alter local nucleosome accessibility to favor a chromatin landscape that is more open at key regulatory genes. In addition to these differentially accessible loci, we also define a trend in an increased genome-wide chromatin accessibility (**Figure 4D**). These accessibility changes are likely to have complex downstream consequences on TF binding and gene expression. Motif analysis of differentially accessible regions identifies enrichment of AP-1/bZIP family motifs, which suggests H3R131C-open loci preferentially facilitates the binding of the AP-1/bZIP families of TFs, which are well-established drivers of oncogenic transcriptional programs including cellular migration, invasion (47). Future studies can further characterize the role of these TF families in mediating the H3R131C-driven oncogenic phenotypes including cellular migration and invasion, and determine whether their occupancy is causally linked to the observed transcriptional and phenotypic changes Given the complex interactions between chromatin accessibility and histone PTMs, future studies examining histone PTM depositions, for example, enhancer-associated H3K27ac or promoter-associated H3K4me3 at differentially accessible regions could define whether H3R131C-driven chromatin opening is sufficient to promote active transcription, or whether secondary remodeling events serve as intermediaries between chromatin accessibility and transcriptional output.

Integration of ATAC-seq and RNA-seq data identified a focused set of genes characterized by concordant increases in both chromatin accessibility and transcript abundance, among which placental growth factor (PGF) represents a compelling candidate mediator of H3R131C-associated oncogenic phenotypes. PGF is a member of the VEGF family that signals through VEGFR1/FLT1 to promote angiogenesis(45), tumor cell survival, and metastatic dissemination (48), and has been associated with poor prognosis in breast cancer (49) and melanoma (50, 51), two tumor types in which genomic alterations encoding H3R131C recurs. The increased accessibility at the PGF promoter in H3R131C-expressing cells, coupled with elevated PGF transcript abundance, raises the possibility that H3R131C-driven nucleosome destabilization at this locus directly licenses PGF expression to support a pro-metastatic tumor microenvironment. Similarly, upregulation of SOX5 — a transcription factor implicated in epithelial-to-mesenchymal transition and invasive behavior in multiple cancers(46) — may contribute to the enhanced migration and invasion that are observes in H3R131C-expressing models. Future functional studies examining the consequences of PGF and SOX5 depletion in the context of H3R131C expression will be important to establish whether these targets are causal drivers of the observed phenotypes.

In summary, here we provide evidence that recurrent genomic alteration that leads to expression of the H3R131C protein represent cancer-associated histone changes that remodel chromatin architecture to drive transcriptional programs associated with migration and invasion, which may support tumor aggressiveness. These findings expand the known repertoire of probable oncohistones beyond PTM-proximal N-terminal tail mutations and highlight the globular domain of histone H3 as a functionally important domain which perturbation can contribute to malignancy. Further characterization of H3R131C *in vivo*, and identification of the transcription factor networks and chromatin regulatory mechanisms downstream of H3R131C-driven accessibility changes, will be essential to understand how this mutation cooperates with co-occurring oncogenic alterations to promote cancer progression. This mechanistic insight may provide insight into therapeutically actionable opportunities for the treatment of cancers characterized by globular domain histone H3 genomic alterations.

## Experimental procedures

### Data mining via COSMIC and cBioPortal

Publicly available patient tumor sequencing data from COSMIC and cBioPortal databases (32, 33) were mined for somatic H3R131C genomic alterations in any of the 15 H3 genes. Co-occurring genomic alterations were defined through specific datamining using the cBioPortal database. From these datasets, cancer pathology, total number of genomic alterations, metastatic presentation, allele frequency, and identity of co-occurring genomic alterations were recorded.

### PyMOL Modeling

PyMOL (Shroedinger) was used for molecular modeling. PDB ID 5X7X (52)was used as the structural template, histone H3 was isolated and the H3R131 residue was identified. The arginine at position 131 was replaced with cysteine using the PyMOL mutagenesis tool. Impacts of H3R131C substitution were assessed by examining the changes in charge and proximity to neighboring residues. Distances between the residue that was mutated and the surrounding amino acids were measured in Å. Images were exported at high resolution.

### Site Directed Mutagenesis and Confirmation

Site directed mutagenesis (SDM) was performed to introduce the H3R131C substitution into the pBabe-puro dH3.3-TY1-IRES-GFP plasmid. Primers for site directed mutagenesis were designed from the web-based Agilent quick change primer design tool. SDM was performed using the manufacturer’s protocol (Agilent quick change II XL) SDM kit and transformed into XL-10 gold ultracompetent cells. Individual colonies were cultured, plasmid DNA extracted (Qiagen), and sanger sequencing was used to confirm the nucleotide changes that encode H3R131C mutation.

Forward Primer: dH3.3 R131C QC F: 5’ – CCC GCT CGC CAC AGA TGC GTC TGG C – 3’. Reverse Primer: dH3.3 R131C QC R: 5’ – GCC AGA CGC ATC TGT GGC GAG CGG G. – 3’.

### Cell lines and culturing

Human mammary epithelial cells (HMEC), HEK293T, and MCF7 cells were STR profiled (ATCC) and maintained at 37°C and 5% CO_2_. HMECs were cultured in DMEM/F12 medium media supplemented with 0.6% FBS, 0.01 µg/ml EGF, 10 µg/ml insulin, 0.025 µg/ml hydrocortisone, 1 ng/ml cholera toxin, 2.5 µg/ml Amphotericin B, and 1% Penicillin-Streptomycin. MCF7 and HEK293T cells were cultured in RPMI or DMEM supplemented with 10% FBS and 1% Penicillin-Streptomycin, respectively. Cells were routinely tested for mycoplasma contamination (Lonza MycoAlert).

### Transfection and transduction

HEK293T cells were transiently transfected with pBabe-puro dH3.3R131C-TY1-IRES-GFP, retroviral packaging gag/pol and VSVg plasmids using the 2X BES calcium phosphate method. 72 hours post transfection, viral supernatant was collected, filtered through a 0.45 µM filter, and concentrated using protein concentrating columns (Prometheus #84-572) by centrifugation at 2500 x g for 15 min at 4°C. Concentrated retrovirus in a volume of less than 500 µL was combined with 10 ug/mL of polybrene and used to transduce HMECs or MCF7 cells. The resulting transduced HMECs and MCF7 cells were and selected using puromycin (1 μg/mL in HMECs, 0.25 μg/mL in MCF7) followed by GFP fluorescence associated cell sorting (FACS) in which the top 50% GFP positive cells were retained for culturing as a pooled population. The selected pooled stable cell lines were then tested to confirm H3R131C-TY1 or H3-TY1 expression by immunoblot.

### Protein extraction and immunoblotting

Histones were extracted using acid extraction as previously described(35, 53, 54). In brief, cells were washed with ice cold PBS, scraped into tubes, and centrifuged. The resulting cell pellets were resuspended in triton extraction buffer ([TEB]; PBS + 0.5% TritonX-100) with 1X protease inhibitors (7.5 μM aprotinin, 0.5 mM leupeptin, 250 μM bestatin, 25 mM AEBSF–HCl). After TEB resuspension and homogenization, 0.8 M HCl was added to solubilize histone proteins and incubated for 20 min on ice and 12000 x g for 10 min. The supernatant was transferred to a clean tube and histones were precipitated with 50% Trichloroacetic Acid. The precipitated histones were collected by centrifugation (12000 x g for 20 min at 4°C) and then washed once with an excess of ice cold acetone-0.3N HCl and then washed twice with ice cold 100% acetone. Pellets were dried at 55°C until translucent. Proteins including histones were resuspended in deionized water and supplemented with Tris-HCl (pH 8.0), 4M NaOH and 1X protease inhibitors. Whole cell protein lysate was extracted on ice using RIPA buffer (50 mM Tris-HCl (pH 7.4), 150 mM NaCl, 1 mM EDTA, 1% NP-40, 1% Na-deoxycholate, 0.1% SDS) with 1X protease inhibitors. Supernatant protein was collected by centrifugation at 15000 x g for 10 min at 4°C. Protein concentration was determined via Bradford assay. Forty ug of acid extracted protein lysate was separated using 16% SDS-PAGE, transferred onto 0.2 µm nitrocellulose membrane at 100V for 1 hr 10 min, and blocked with 5% nonfat milk in TBST for 1 hr. Membranes were incubated with primary antibodies overnight at 4C and secondary antibodies and then developed using the LICOR Odyssey Clx. Primary antibodies and dilutions used are as follows: TY1 (Diagenode C15200054; 1:1000), H3.3 (Abcam ab17899; 1:1000), H3 (Abcam ab1791; 1:10000). Secondary antibodies used include fluorophore-conjugated goat anti-rabbit 680 IgG (LiCor, 925–68 071; 1:10,000) and goat anti-mouse 800 IgG (LiCor, 926–32 210; 1:10,000).

### In vitro proliferation assay

Subconfluent cells were trypsinized, counted, and seeded at 3,000 cells per well in a 96-well plate in full growth media in triplicate. At 24, 48, 72, and 96 hrs post seeding, plates were fixed (50% MeOH + 10% acetic acid) for 15 min, stained with crystal violet (0.2% crystal violet + 10% EtOH) for 15 min, washed in deionized water until clear, and dried. Plates were destained with destain solution (40% EtOH and 10% acetic acid) for 15 min and absorbance was measured at 595 nm. Proliferation curves were created after averaging three biological replicates and normalizing them into cells expressing WT H3.

### Clonogenic assay

Sub confluent cells were trypsinized, counted, and 500 cells per well were seeded in duplicate in a 6-well plate in technical duplicate. Media was changed every three days, and 14 days after initiating the experiment. the cells were fixed with a fixative buffer (50% MeOH + 10% acetic acid) for 15 min, stained with crystal violet (0.2% crystal violet + 10% EtOH) for 15 min, and washed in deionized water until clear. The plates were dried and imaged using a scanner. The crystal violet was destained using a destaining solution (40% EtOH and 10% acetic acid). Absorbance was measured at 595 nm and the colony forming potential of H3R131C was compared relative to WT H3.

### Boyden chamber migration assay

Sub confluent cells were trypsinized, resuspended in serum free media, counted, and 1 x 10^5^ cells were seeded in duplicate in the boyden chamber insert in a 24-well plate (Corning). 600 μL of complete media was added in the bottom chamber to serve as a chemoattractant. After 8 hr, the non-migrated cells were removed from the upper surface of the boyden chamber using a Qtip®. Migrated cells at the bottom of the boyden chamber were fixed in 70% EtOH for 10 min, stained with crystal violet (0.2% crystal violet + 10% EtOH) for 15 min, washed in deionized water until clear, and air dried. The membranes were cut, mounted on slides, and imaged using a light microscope (Leica Delimited). Migrated cells were quantified using ImageJ. Migration potential of H3R131C was compared to WT H3.

### Boyden chamber invasion assay

On ice, matrigel (Corning, 254234) was diluted to 1 mg/mL in serum free media and 100 μL was applied to the surface of the chamber to solidify. Subconfluent cells were trypsinized, resuspended in serum free media, counted, and 1 x 10^5^ cells were seeded in technical duplicate on top of the Matrigel layer. Complete medium in the lower chamber was used to serve as a chemoattractant. After 48 hrs, the non-invading cells and matrigel were removed from the chamber using a Qtip®. Invaded cells were fixed with 70% EtOH for 10 min, stained with crystal violet (0.2% crystal violet + 10% EtOH) for 15 min, washed in deionized water until clear, air dried, and cut and mounted onto slides. The slides were imaged and invaded cells were quantified using ImageJ. Invasion potential of H3R131C was compared relative to WT H3.

### RNA isolation and sequencing

RNA was isolated from HMECDD dH3.3R131C-TY1 and dH3.3-TY1 cells with on column DNaseI digestion (Qiagen RNeasy). RNA libraries were prepared in house (Tecan Magic Prep) and sequenced via the Illumina platform with 150bp paired end (PE) reads at a depth of 30M PE reads (Admera Health LLC).

### ATAC sample preparation and sequencing

For ATAC sequencing, chromatin accessibility libraries were prepared using a standard Omni-ATAC protocol as previously described (35, 41). In brief, cells were counted and 50,000 cells were washed with cold ATAC-seq resuspension buffer (RSB; 10 mM Tris-HCl pH 7.4, 10 mM NaCl, and 3 mM MgCl2 in water), and permeabilized with ATAC-seq lysis buffer (RSB supplemented with 0.1% NP40, 0.1% Tween-20, and 0.01% digitonin). For the transposition step, cells were resuspended in 44 µl of transposition mix (25 µl 2× TD buffer, 2.5 µl transposase, 16.5 µl PBS, 0.5 µl 1% digitonin, and 0.5 µl 10% Tween-20) and incubated at 37°C for 40 min in a thermomixer with shaking at 1,000 RPM. The DNA fragments were cleaned up with the DNA Clean & Concentrator kit (Zymo Research, D4014) and PCR amplified for 8-12 cycles with Illumina Nextera adaptors using Kapa SYBR® FAST (Roche, 07959427001). The number of PCR cycles was determined based on monitoring for a change in fluorescence of 100,000 or a ΔRn of 0.1, after which the PCR was stopped and the samples removed. After amplification, a second cleanup with 1.8x Ampure XP beads (Aline biosciences, Cat. C-1003-50) was performed and library eluted in 10 mM Tris-HCl. Fragment size distribution and concentration were evaluated via Bioanalyzer High Sensitivity DNA kit (Agilent, 5067-4626), and libraries were sequenced to a target depth of 75M paired-end reads (Admera Health LLC).

### Next generation sequencing (NGS) analysis

Post RNA sequencing, the resulting FASTQ files were aligned to the human reference genome (build GRChg38) and transcript abundance was quantified using salmon(55) (Illumina DRAGEN v4.3.6). The DESeq2 (56) package was used to identify differentially expressed transcripts between the H3R131C and WT H3 HMECDD cells. Transcripts with adjusted *P*-value ≤ .05 and log_2_(FoldChange)| ≥ ±1 were considered differentially expressed. EnhancedVolcano was used to generate volcano plots. Fast gene set enrichment analysis (FGSEA) of the pre-ranked gene list was performed using clusterProfiler (57-59) and fgsea, using the msigdb hallmark gene set (60, 61). Gene ontology (GO) enrichment analysis of differentially expressed genes was performed using clusterProfiler to identify enriched biological processes among upregulated and downregulated gene sets. Normalized read counts for select target genes were extracted from DESeq2 output and plotted in R Post ATAC sequencing, the resulting FASTQ files were processed as follows: Raw reads were processed using Illumina DRAGEN v4.3.6. Mitochondrial reads, duplicates, non-unique alignment reads, and black-listed reads were removed using Samtools (62) and bedtools (63). The final .bam files were then indexed and peaks were called using MACS3 (64). Graphable matrix files were generated using Deeptools (65). ComputeMatrix and heat maps generated via plotHeatmap. Differential accessibility analysis was performed using DIFFbind. BigWig files were then generated using bamCoverage (65). WT H and H3R131C samples were compared to identify regions that gain or lost accessibility and PCA and volcano plots were generated. Nearest genes were annotated with CHIPseeker (66). Motif enrichment analysis was performed using HOMER (42) and MEME Suite (67), and results were integrated to identify enriched motifs. Concordance between chromatin accessibility and gene expression was evaluated using Pearson correlation and visualized using ggplot2 and Venn diagrams.

## Statistical analysis

All the experiments described herein were performed in three independent experiments unless otherwise noted. Mean ± SD were calculated and used to plot graphs unless otherwise noted. Statistical significance (*P* ≤ .05) of differences between WT H3 and H3R131C groups was determined by Student’s *t*-test. To compare average allele frequency and aneuploidy scores amongst alterations of H3R131, H3K27, and H3K36, a one-way ANOVA was performed using GraphPad Prism. To analyze the probability of cancer diagnosis associated with alterations in H3 residues, the gnomAD (34) database was surveyed for alterations in all H3 encoding genes for variants that recur in an undiagnosed population. The number of altered alleles was determined and compared with the number of alterations of H3 residues found in cBio and COSMIC databases using a logistic regression model and visualized in R.

## Acknowledgements

The authors would like to thank the members of the Spangle, Corbett, and Hong labs, including Team Histone, for their research discussions and support. We thank Dr Dan Yan, Department of Pediatrics, Emory University for providing support for the Sartorius Incucyte and Emory pediatric flow core.

## Author Contributions

SVL, KS, AJK, EJH, and JMS designed experiments, SVL, KS, AJK, and EJH conducted experiments. SVL, KS, and AJK analyzed data. SAS, AHC, and JMS oversaw the studies and provided directional input. SVL wrote the first draft of the manuscript with input from KS, AJK, AHC and JMS. JMS was responsible for broad project direction and funding acquisition. All authors reviewed and edited the manuscript.

## Data Availability

The data generated in this study are publicly available in the Gene Expression Omnibus at GSA (RNA and ATAC, GEO accession numbers pending). Cell lines and plasmids are available upon request.

