## Supplementary data for "Globular domain histone H3R131C mutation remodels chromatin accessibility to promote oncogenic transcriptional programs"

### Supporting information

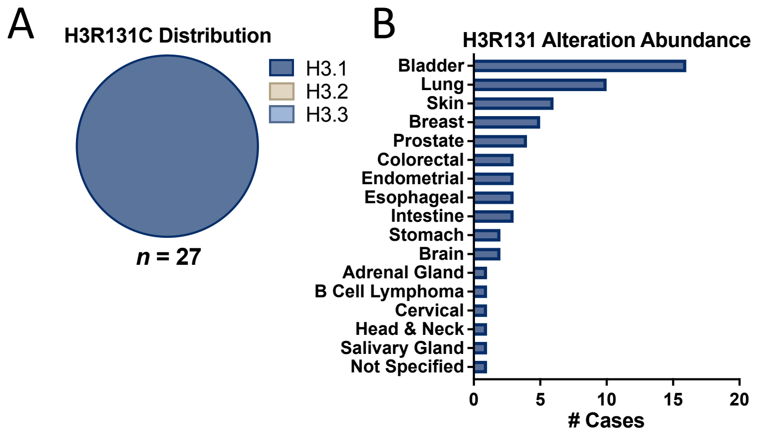


**Supplementary Figure 1: H3R131 genomic alterations recur in human cancers.**
Data collected from cBioPortal and COSMIC databases. A) Distribution of genomic alterations in histone H3 genes that encode H3R131C. B) Distribution of identified cancer diagnoses that harbor changes in H3R131.

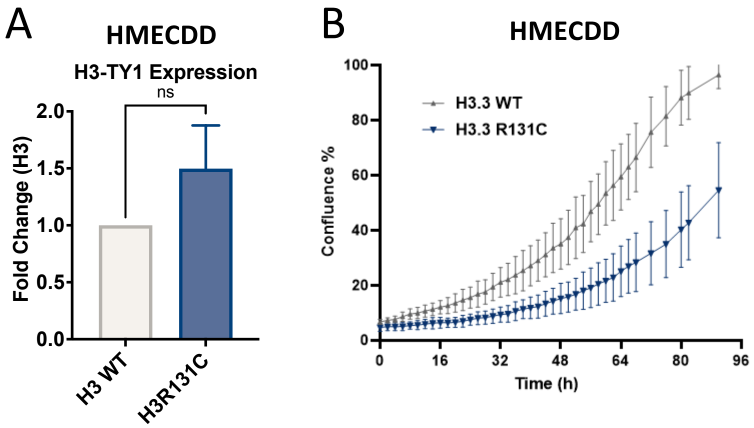


**Supplementary Figure 2: H3R131C expression supports oncogenic phenotypes.**
A) Quantification of Figure 3A: HMECDD cells stably transduced with pBabePuro H3.3-TY1 or H3.3R131C and lysates acid extracted. *n = 3*. B) Stable HMECDD cells expressing the indicated H3.3 mutant proteins were seeded and cell proliferation measured over the indicated time course using the Incucyte platform, *n = 2*.

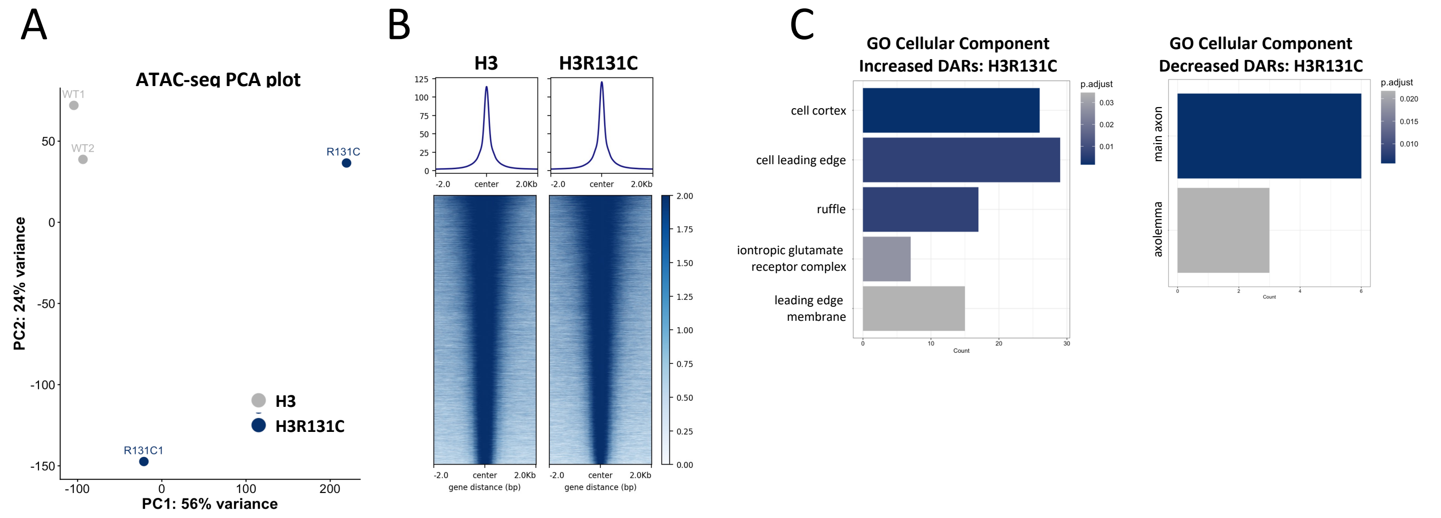


**Supplementary Figure 3: H3R131C alters chromatin accessibility.** A) Principal component analysis (PCA) plot derived from ATAC-seq data from HMECDD cells stably expressing H3R131C-TY1 or H3-TY1. *n = 2*. B) Tornado plots assessing chromatin accessibility in HMECDD cells stably transduced with H3 or H3R131C. Plots represent the top 15,000 peaks identified in WT H3.3 and queried in H3.3 R131C. Representative tornado plot shown, *n = 2*. C) GO depicting upregulated and downregulated cellular components from HMECDD H3R131C-TY1 cells compared to HMECDD H3-TY1 cells.

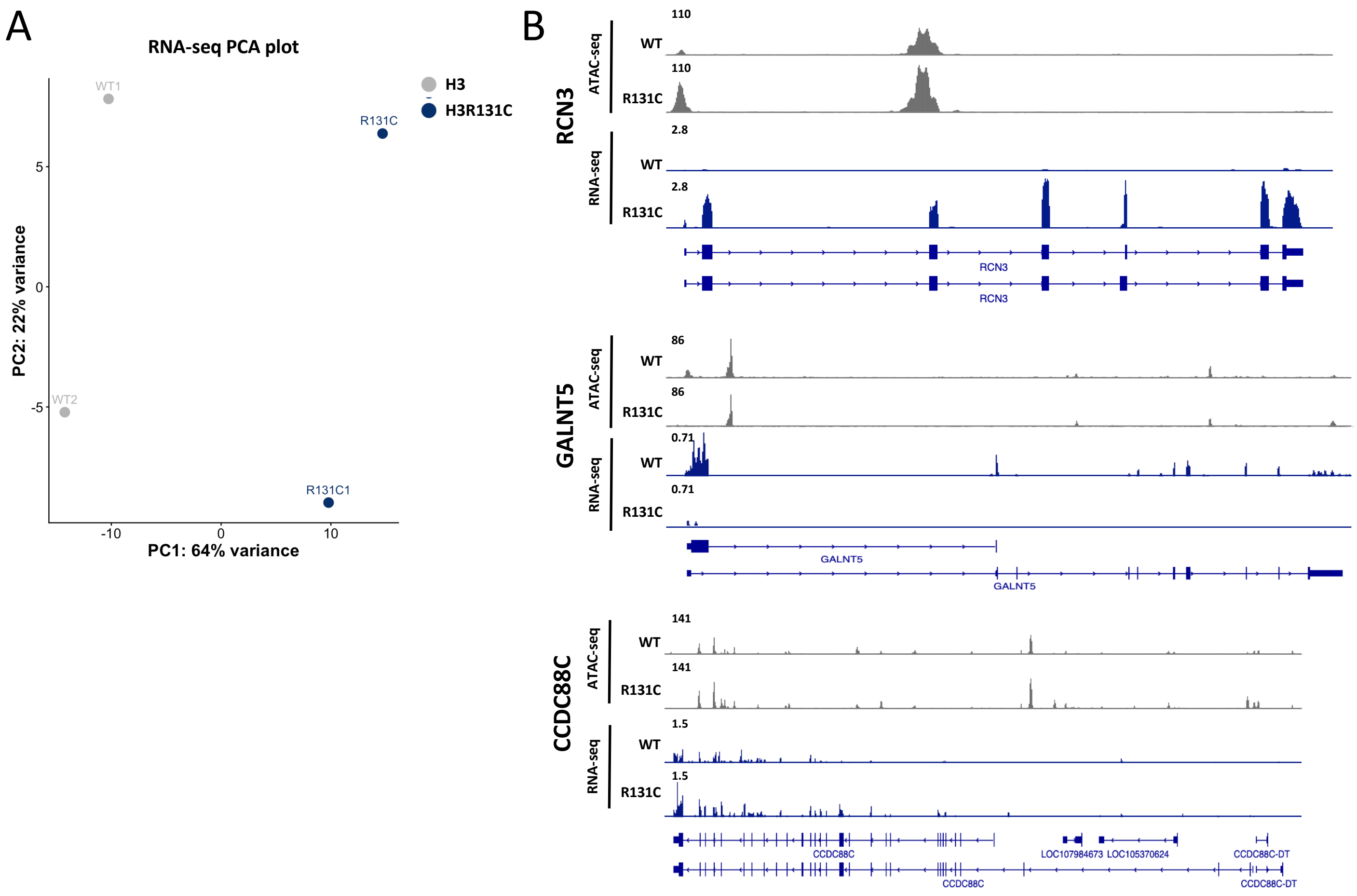


**Supplementary Figure 4. H3R131C chromatin remodeling drives transcriptional changes that support H3R131C oncogenic phenotypes**. A) Principal component analysis (PCA) plot derived from RNA-seq data from HMECDD cells stably expressing H3R131C-TY1 or H3-TY1. *n = 2*. B) IGV tracks from ATAC-seq and RNA-seq of selected genes.
